# RAPPL-Seq Enables Rapid, High-Sensitivity Ribosome Profiling of Scarce Samples

**DOI:** 10.64898/2026.10.20.760540

**Authors:** Ajin Park, Krzysztof J. Szkop, Chaehee Lim, Jessey Erath, Jiayi Huang, Dimitrios Mathios, Zhiyu Dai, Xueyan He, Slavica Pavlovic-Djuranovic, Ola Larsson, Ivan Topisirovic, Sergej Djuranovic, Chun-Kan Chen

## Abstract

Ribosome profiling is a powerful method for transcriptome-wide monitoring of translation activity *in vivo*, but its use is limited by lengthy protocols, reliance on specialized equipment and inefficient recovery of ribosome-protected fragments (RPFs). To overcome these challenges, we developed RAPPL-Seq, a magnetic bead-based affinity capture method that enables fast and efficient RPF isolation. RAPPL-Seq achieves high recovery efficiency, short processing time, and robust performance with limited input material. It is broadly applicable across multiple sample types, including cell lines, primary tissues, and human biospecimens. RAPPL-Seq can also be paired with translation inhibitors to characterize inhibitor- and tissue-specific mRNA translation. RAPPL-Seq efficiently captured translatomes of glioblastoma (GBM) and adjacent non-tumor brain tissues from patient biopsies. Together, these findings establish RAPPL-Seq as a straightforward, easily accessible, and highly sensitive approach that extends translatome studies to scarce experimental and clinical samples, opening new avenues for mechanistic insight and therapeutic discovery.

## INTRODUCTION

Ribosome profiling is a powerful high-throughput technique that enables transcriptome-wide monitoring of ribosome occupancy *in vivo* through deep sequencing^1–3^. This approach provides insights into alterations in mRNA translation across baseline, disease, and stress conditions, thereby revealing regulatory elements and mechanisms of translational control^4–7^. The method relies on generating ribosome-protected fragments (RPFs) through ribonuclease digestion, isolating as well as sequencing of these fragments, to obtain ribosome footprints on RNAs^1–3^. These footprints allow quantitative assessment of translation activity and ribosome occupancy across transcripts on a genome-wide scale^3^. In addition, RPFs can uncover translons outside canonical annotations, such as those embedded within 5’ untranslated regions (5’ UTRs), enabling investigation of the regulatory and functional roles of non-canonical ORFs under stress or disease conditions^4,6,8,9^. Ribosome profiling is also broadly applicable across organisms, including prokaryotes, where, for example, it can be used to monitor bacterial translation in response to antibiotic treatment and facilitate antibiotic screening or mechanistic studies^10–14^.

Translational reprogramming plays a pivotal role in cancer and other diseases including neurological and metabolic disorders^15–18^. For example, tumors dynamically remodel the translatome to rapidly reshape the proteome and drive tumor initiation, progression and dissemination^15–18^. Capturing changes in the translatome between non-transformed and malignant cells as well as between malignant cells at different steps of tumor evolution is therefore of paramount importance for uncovering molecular underpinnings and vulnerabilities of neoplasia. In this context, the inherent scarcity of patient biopsy material hinders approaches based on the traditional ribosome profiling, which require substantial starting material. Consequently, most translatome studies in cancer and other pathological states rely heavily on cell culture models, such as cell lines or organoid cultures^19,20^. However, these systems may not accurately recapitulate the complex features of neoplasia *in vivo* within its natural tumor microenvironment (TME) and under physiological levels of nutrients growth factors and hormones. Therefore, developing a highly sensitive translatome analysis technology compatible with scarce clinical biospecimens is crucial to establish the role of the translational apparatus in the context of cancer and other diseases.

Traditionally, ribosome profiling relies on isolating RPFs using sucrose cushions or gradients^2,21–23^. This requires relatively large amounts of input material in part due to substantial sample loss during centrifugation on sucrose cushions or gradients and RNA precipitation^24^. As a result, the traditional Ribo-seq approach is not suitable for clinical samples, such as patient biopsies and primary tissues, where input material is often limited. In addition, the lengthy sucrose cushion and/or gradient procedures can lead to RNA degradation, particularly affecting transcripts that are expressed or translated at low levels. Finally, these methods require specialized ultracentrifugation and fractionation equipment, which may not be readily accessible. On the other hand, immunoprecipitation (IP)-based ribosome affinity purification approaches^25,26^ can bypass sucrose cushion or gradient methods but are limited by several drawbacks: these methods often require introducing exogenous tag-fused translation factors, which makes them unsuitable for primary tissues or clinical samples. In addition, IP-based pull-downs typically suffer from low efficiency, resulting in substantial loss of input material and limiting their applicability to low-input samples. Together, these challenges underscore the need for a rapid, streamlined RPF isolation method that is efficient, robust, broadly accessible, and compatible with low-input samples.

Recently, our group developed RNA Affinity Purification using Poly-Lysine (RAPPL), a method that isolates ribosomes and their associated RNA fragments through electrostatic interactions between the negatively charged RNA backbone and the positively charged amino groups of poly-lysine on magnetic beads^24^. By bypassing sucrose gradients/cushions and RNA precipitation, RAPPL enables rapid and efficient capture of ribosomes and associated RNA fragments, making it well suited for isolating RPFs from a variety of samples of different origins. Building on this approach, we adapted RAPPL for translatome profiling and established RAPPL-Seq, a technique that allows robust, transcriptome-wide monitoring of ribosome occupancy from limited material including human tissue specimens. RAPPL-Seq is compatible with translation inhibitor treatments and broadly applicable across multiple sample types, including cell lines, primary tissues, and biopsy specimens. Finally, by applying RAPPL-Seq, we captured the translatomes of glioblastoma (GBM) and unaffected neighboring brain tissue from patient biopsy. This demonstrates RAPPL-Seq is capable of reliably capturing translatome profiles in research and clinical settings where sample scarcity is a primary constraint.

## RESULTS

### RAPPL-Seq Efficiently Recovers and Enriches Ribosome-Protected Fragments

To rapidly and efficiently capture ribosome-protected fragments (RPFs) for subsequent ribosome profiling, we developed RAPPL-Seq, which was adapted from RAPPL - a simple, rapid, and cost-effective method for isolating ribosomes^24^. Briefly, cells were flash-frozen in liquid nitrogen (LN_2_) to immobilize ribosomes on RNA, followed by lysis in the presence of cycloheximide (CHX) and RNase I treatment to generate RNA fragments protected of nuclease digestion by ribosomes. Ribosomes and their associated RNA fragments were then captured using poly-lysine magnetic beads via electrostatic interactions between the negatively charged RNA backbone and the positively charged amino groups of poly-lysine^24^ **(Figure 1a and Supplemental Figure 1a)**. After washing to remove unbound material, RPFs were eluted using poly-glutamate and purified by TBE-Urea gel extraction to isolate RNA fragments of the desired size (25-35 nt for RPFs^24^ ; **Figure 1b**). The purified RPFs were then subjected to rRNA and tRNA depletion, followed by standard ribosome profiling library preparation (LACEseq)^27,28^ and next-generation sequencing **(Figure 1a; Methods)**.

**Figure 1.**
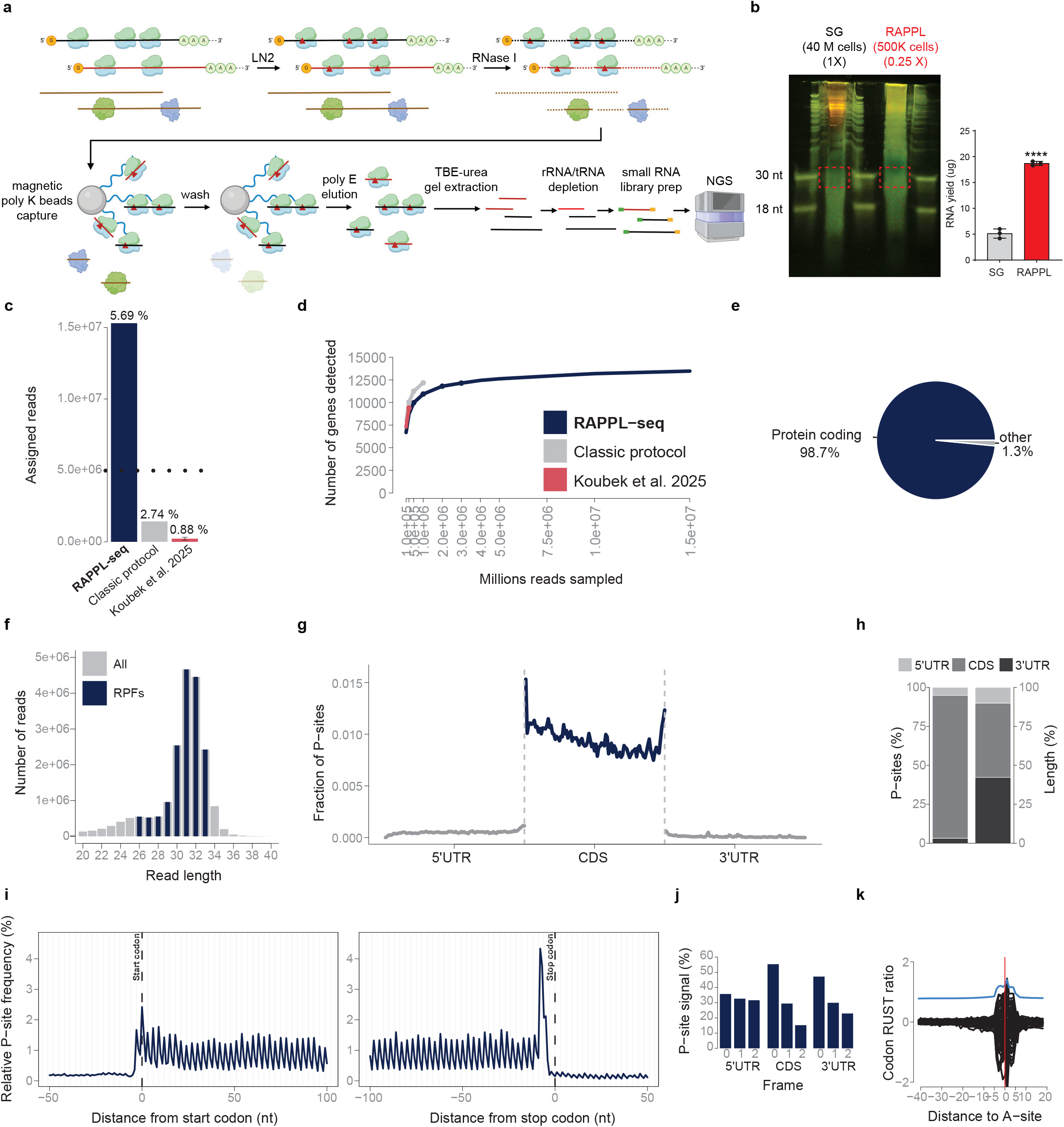
RAPPL-Seq enables rapid and efficient capture of RPFs. **(a)** Schematic overview of RAPPL-Seq, including RPF capture and purification, library preparation, and sequencing. **(b)** Comparison of RNA recovery yields from 40,000,000 HEK293T cells processed by sucrose gradient (SG) versus 500,000 HEK293T cells processed by RAPPL-Seq (RAPPL) (25% of total recovered RNAs), visualized by TBE-Urea gel electrophoresis (left) and quantified by bar graph (right). Bar graphs represent mean ± SD from three independent replicates. **** p < 0.0001 by two-sample t-test compared with SG. Red dashed boxes indicate the regions extracted for enriching for 25-35 nt RPFs. **(c)** Assigned RPF read counts and percentages out of total number of reads and **(d)** Library complexity curves showing the number of genes detected across sampled sequencing depths for RAPPL-Seq, the classic protocol and the protocol reported by Koubek et al.^92^. The dotted line in (c) indicates an approximate benchmark for the differential translation analysis^29^. **(e)** Distribution of RNA biotypes among RAPPL-Seq reads mapped to GENECODE v50 transcripts. **(f)** Read-length distributions for all reads (grey) and RPFs (blue). **(g)** Metagene profile of pooled P-site counts across the 5’UTR, CDS and 3’UTR, with each region divided into 100 equal sized bins. **(h)** Percentage of P-sites assigned to the 5’ UTR, CDS, and 3’ UTR in RAPPL-Seq libraries (left), compared to the expected distribution under random RNA fragmentation (right). **(i)** Metagene profiles showing relative P-site frequencies around annotated start codons (left) and stop codons (right), highlighting triplet periodicity within the CDS. **(j)** Distribution of P-sites among reading frames 0, 1, and 2 within the 5′ UTR, CDS, and 3′ UTR. **(k)** Codon RUST ratios (log_2_) plotted relative to the A-site, indicated by the red vertical line. The corresponding Kullback–Leibler divergence is shown in blue.

We first utilized 1,000,000 HEK293T cells to benchmark RAPPL-Seq, as different ribosome profiling approaches have been extensively characterized in this cell model. Compared with the traditional sucrose gradient-based purification method, RAPPL-Seq recovered ∼16-fold more RNA from 20-fold fewer cells **(Figure 1b)**. Notably, the observed higher recovery of RNA using poly-lysine magnetic beads resulted in a higher percentage of translatome assigned reads (1.5×10^7^ and 5.69% of total reads) compared to recently published (less than a 1×10^6^ and 0.88%) and classical (1.5×10^6^ and 2.74%) Ribo-seq protocols **(Figure 1c)**. Importantly, the high RNA recovery obtained with RAPPL (∼80% in a single elution) was not due to the capture of non-ribosome-associated RNAs, as these RNA species constituted a smaller portion of the total yield, comparable to levels seen in current ribosome profiling protocols **(Supplemental Figure 1b and 1c)**. Moreover, RAPPL recovered ∼90% of ribosomes from pre-isolated sucrose gradient fractions, thus suggesting that the enhanced recovery efficiency results from the inherent high-affinity interaction between poly-lysine and RNA-enriched complexes **(Supplemental Figure 1b and 1c)**^24^. These results indicate that RAPPL has superior ribosome capture efficiency compared with current ribosome profiling methods.

The proportion of assigned RPF reads (∼6% of total reads) was substantially higher than that reported for previously published methods, enabling the recovery of sufficient reads for differential translation analysis at a relatively modest sequencing depth^29^ **(Figure 1c)**. RAPPL-Seq showed high library complexity and approached saturation in gene detection compared with other ribosome profiling methods **(Figure 1d)**. Furthermore, 98.7% of assigned reads localized specifically within annotated protein-coding sequences **(Figure 1e)**. 1.3% of assigned reads mapped to a subset of long non-coding RNAs (lncRNAs)^30–34^. Analysis of the footprint length distribution of RPFs purified by RAPPL-Seq revealed focused distribution of the RPFs between 29 and 33 nt with highest number of assigned reads distributed at 31 and 32 nt length **(Figure 1f)**. This is consistent with previous studies and corresponds to the size of the RNA fragments protected by translating ribosomes^7,35,36^. RAPPL-Seq reads, aligned to the P-site, were distributed mainly within the CDS, with a relatively small number of reads in the 5’ and 3’UTRs **(Figure 1g and 1h).** The captured RPFs demonstrated clear triplet periodicity along the CDS and ribosome pausing at translation initiation sites (start codon) and translation termination sites (stop codon)^2,3,37^ **(Figure 1i)**. The CDS occupancy indicated a preference for in-frame RPFs (+0 frame) among the footprint fragments, thereby illustrating predominant capture of ribosome footprints in the correct reading frame **(Figure 1j)**. A similar, albeit less pronounced, distribution was observed in the 3′ UTR but not the 5′ UTR, suggesting possible stop codon readthrough events **(Figure 1j)**. Finally, we observed similar codon RUST ratios **(Figure 1k)** matching previous studies and indicating the absence of potential biases related to other sequence determinants^1,3,37,38^. Together, RAPPL-Seq provides a simple, rapid, and robust approach for efficiently enriching RPFs from cultured cells, enabling high-quality ribosome profiling.

### RAPPL-Seq Captures Effects of Inhibitors Acting at Different Phases of Translation

Translation inhibitors can be used to stall ribosomes in defined conformations, enabling the capture of RPFs at distinct stages of translation thereby providing important insights into translational regulation^1,4,39–42^. In addition to the above mentioned LN_2_ flash-freezing for ribosome immobilization, RAPPL-Seq is readily compatible with inhibitor treatments. Cycloheximide (CHX) blocks ribosomal translocation, resulting in global ribosome stalling on mRNAs^4,41,42^, whereas harringtonine (Harr) prevents elongation after initiation, causing ribosomes to accumulate at initiation codons and thereby allowing the mapping of translation initiation sites (TISs)^4,41,42^.

To benchmark inhibitor treatments, 500,000 HEK293T cells were pre-incubated with CHX or Harr at 37 °C for 10 minutes prior to standard RAPPL-Seq processing. Both CHX and Harr treatments yielded ribosome footprint sizes and frame distributions comparable to those observed following LN_2_ flash-freezing **(Figure 2a and Supplementary Figure 2a**), but P-site reads were enriched in the 5’UTR with Harr and, albeit to a lesser extent, for CHX treatment **(Figure 2b-c)**. The elevated 5’UTR signal observed following CHX treatment likely reflects a previously identified technical artifact of CHX pretreatment on ribosome distribution near TISs^43^. CHX treatment showed overall metagene profiles of ribosomal periodicity (**Figure 2d, blue and green lines)** and codon usage frequencies **(Supplemental Figure 2b)** comparable to those observed in LN_2_ flash-frozen cells, with a clear signal at start and stop codons. As expected, Harr treatment resulted in reduced ribosome occupancy across most codons compared with CHX-treated and LN2 flash-frozen cells, consistent with a block of elongation at the start codon **(Supplemental Figure 2a)**^4,41,42^. Notably, after Harr treatment, RPFs were enriched in 5’ untranslated regions (5’ UTRs) with a proportion higher than the inherent representation of 5’ UTRs in mRNAs **(Figure 2a-d and Supplementary Figure 2a and 2b)**, which aligns with previous reports of pervasive alternative translation initiation and translation activity in 5’ UTRs of the human genome^6,42,44–46^. Importantly, this enhanced resolution at initiation sites following Harr treatment allowed RAPPL-Seq to identify non-canonical TISs, consistent with previous reports of alternative translation initiation within 5’UTRs^6,42,44–46^. These included upstream (uTIS) and downstream (dTIS) sites relative to annotated TISs (aTISs). This is in agreement with prior ribosome profiling studies using Harr or lactimidomycin as translation inhibitors blocking the first step of elongation^4,41,42,47^. We observed examples of uTIS, dTIS, and uORFs in multiple transcripts including recently reported ZBTB8OS and CMPK1, as well as in more complexed and alternatively spliced transcript of CEBPB transcription factor **(Figure 2e-g)**^48–50^. The uORF-associated reads were also observed in CHX treatment and to a much lesser extent following LN_2_ treatment. We also observed uORFs in RNF10 and RAB6A (not yet been annotated) transcripts, but not in ACTB and CCDC124 **(Supplemental Figure 2c-f)**, indicating the utility of RAPPL-Seq in characterizing alternative translation initiation events. Together, these results highlight the versatility of RAPPL-Seq in integrating multiple translation-stalling strategies, broadening its applicability for dissecting translational regulation and uncovering alternative initiation mechanisms with high resolution.

**Figure 2.**
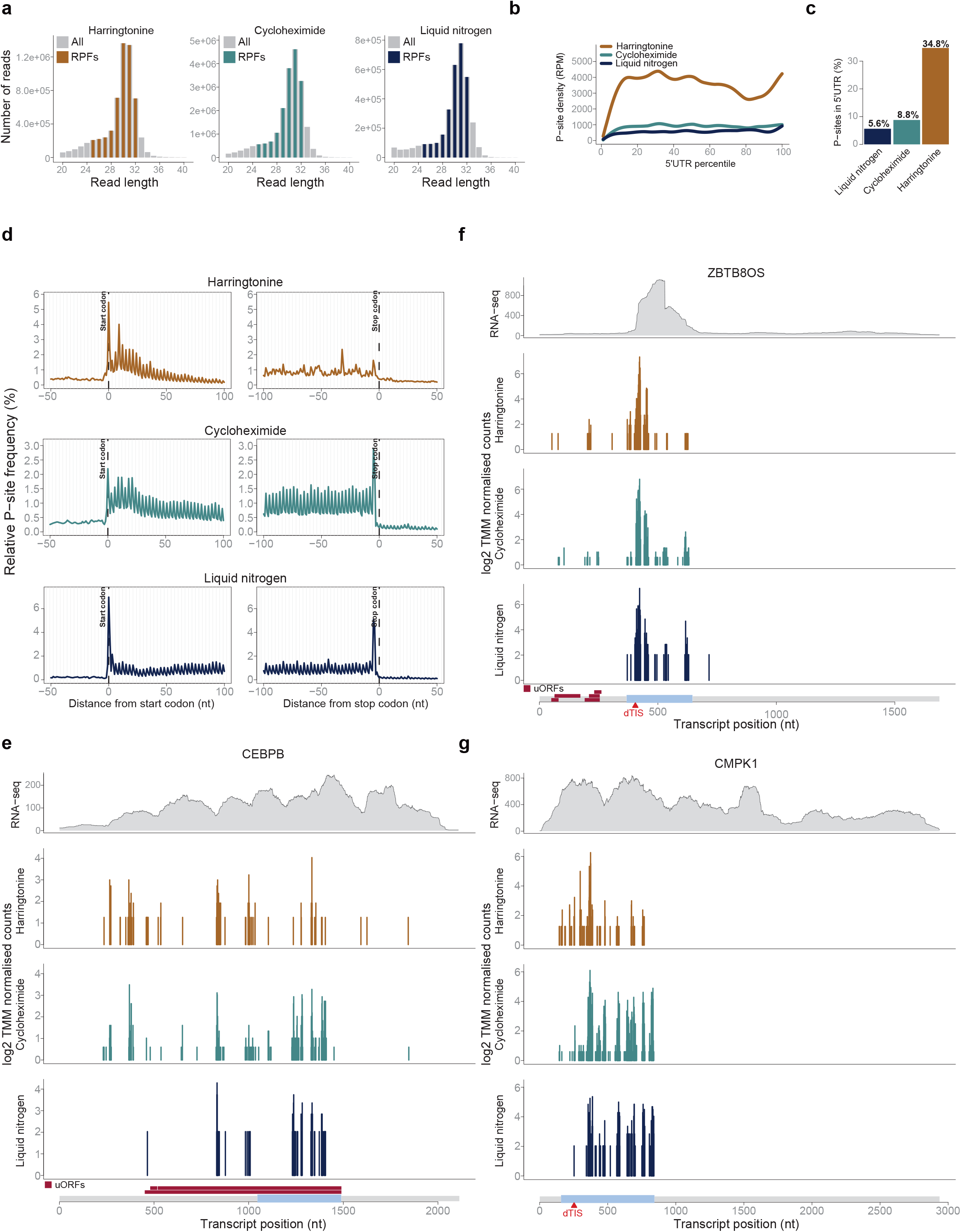
RAPPL-Seq Enables Analysis of Translation Inhibitors Acting at Different Stages. The same colour coding identifies the three conditions throughout the figure: harringtonine (Harr, brown) and cycloheximide (CHX, teal) pretreatments, and flash-freezing in liquid nitrogen (LN_2,_ blue). **(a)** Read length distributions for all reads (grey) and RPFs. **(b)** Metagene profiles of P-site density (reads per million) across the 5’ UTR, with positions expressed as percentiles of UTR length. **(c)** Percentage of P-sites mapping to the 5’UTR for each condition. **(d)** Metagene profiles showing relative P-site frequencies around annotated start codons (left) and stop codons (right), illustrating treatment-dependent differences in triplet periodicity within the CDS. **(e-g)** Representative RNA-seq coverage tracks (raw counts, top) and RAPPL-Seq profiles (log_2_ TMM-normalised counts) under the three conditions for CEBPB **(e)**, ZBTB8OS **(f)**, and CMPK1 **(g)**. Transcript models are shown below each set of profiles.

### RAPPL-Seq Allows Translatome Analysis from Low-Input Samples and Capturing Lowly Expressed Genes

Inefficient recovery of RPFs after ribosome isolation represents a major limitation of current ribosome profiling techniques, particularly for the analysis of small samples with limited cell numbers and the detection of lowly expressed genes **(Supplemental Figure 1b)**^24^. By contrast, the high ribosome capture efficiency of RAPPL-Seq makes it well suited for low-input applications while maintaining good coverage of the translatome. Using 500,000 cells, we obtained robust RPF coverage across the translatome under different inhibitor conditions (**Figure 2 and Supplemental Figure 2)**. To systematically determine the lower input limit of RAPPL-Seq, we performed titrations ranging from 1,000,000 down to 100,000 cells. Although the proportion of assigned reads decreased from the 1,000,000 cells sample, it remained similar between 500,000 and 200,000 cells, with a pronounced decrease observed only at 100,000 cells **(Figure 3a)**. RAPPL-Seq maintained comparable library complexity with inputs as low as 200,000 cells **(Figure 3b)**. Under lower-input conditions (200,000 and 100,000 cells), we also observed a shift in RPF length toward smaller fragment sizes, although this effect was substantially more pronounced at 100,000 **(Figure 3c)**. This shift is likely attributable to RNase I over-digestion, a well-documented obstacle when scaling down input material for Ribo-Seq^35^. Despite the reduction in input, RPFs from 200,000 cells remained predominantly localized to the CDS **(Figure 3d)**. Consistent with this, clear reading frame preference and triplet periodicity were maintained down to 200,000, albeit with modest deterioration in RPF length **(Figure 3e-f)**. Codon usage profiles were also broadly similar **(Figure 3g)**. Similarly, codon RUST profiles remained broadly consistent down even at 100,000 cells, further supporting preservation of RPF characteristics at reduced input **(Figure 3h)**. Strong correlations with the 1,000,000-cell sample supported reproducibility down to 200,000 cells **(Figure 3i-k)**. Further comparison of transcripts with high, moderate, and low RPF **(Supplemental Figure 3a)** indicated similar coverage profiles across multiple examples, including highly expressed DDX1, medium-expressed DDX59, and low-expressed PNISR among other analyzed genes **(Supplemental Figure 3b-d)**. Together, these results demonstrate that RAPPL-Seq can produce robust and high-quality ribosome profiling data from as few as 200,000 cells in this study, thereby significantly extending its applicability to low-input samples^51,52^.

**Figure 3.**
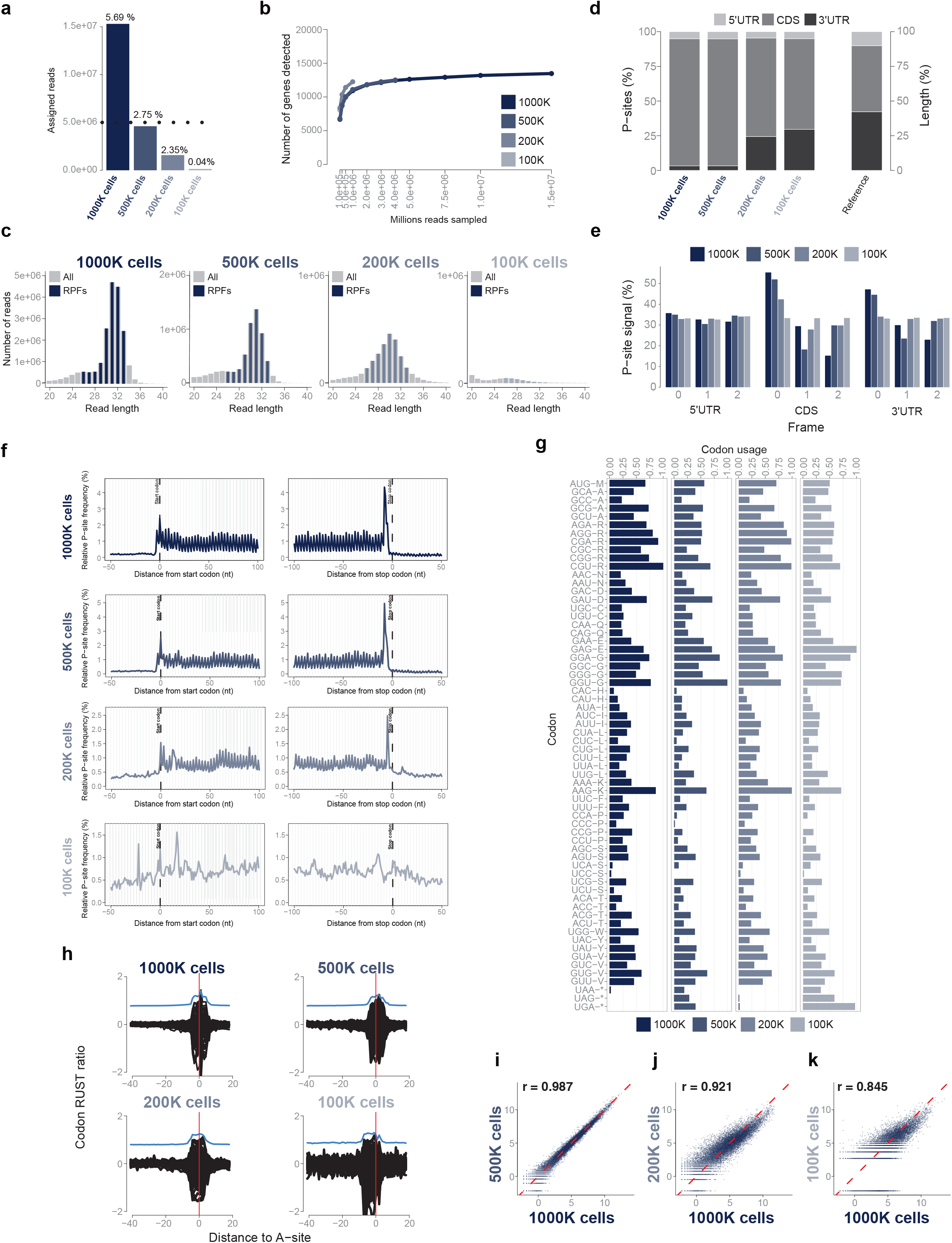
RAPPL-Seq Enables Ribosome Profiling from Low-Input Samples. The same colour coding identifies the four input amounts throughout the figure: 1000K (dark blue), 500K (slate blue), 200K (blue grey), and 100K (light grey) HEK293T cells. **(a)** Assigned RPF read counts and percentages of total reads and **(b)** library complexity curves showing the number of genes detected across sampled sequencing depths for each input amount. The dotted line in panel (a) indicates an approximate benchmark for differential translation analysis^29^. **(c)** Read length distributions for all reads (grey) and RPFs (blue). **(d)** Percentage of P-sites assigned to the 5’ UTR, CDS, and 3’ UTR at each input amount (left), compared to the expected distribution under random RNA fragmentation (right). **(e)** Distribution of P-sites among reading frames 0, 1, and 2 within the 5′ UTR, CDS, and 3′ UTR. **(f)** Metagene profiles showing relative P-site frequencies around annotated start codons (left) and stop codons (right), illustrating input-dependent differences in triplet periodicity within the CDS. **(g)** Codon usage profiles for each input amount. **(h)** Codon RUST ratios (log_2_) plotted relative to the A-site, indicated by the red vertical line. The corresponding Kullback–Leibler divergence is shown in blue. **(i-k)** Reproducibility of RPF abundance across input amounts, assessed by pairwise comparisons of TMM-normalised log_2_ counts per million between 1000K cell dataset and the 500K **(i)**, 200K **(j)**, and 100K **(k)** datasets. Pearson’s correlation coefficients are shown.

### RAPPL-Seq is Broadly Applicable to Human Biopsies and Surgical Specimens

Despite its potential to reveal disease-associated changes in mRNA translation, ribosome profiling has been underutilized for clinical biospecimens, partly due to the scarcity of material in primary tissues and biopsies, as well as loss of material during sucrose cushion and gradient approaches. Having established that RAPPL-Seq enables robust profiling from as few as ∼200,000 HEK293T cells, we next assessed its applicability to primary tissue and clinical biospecimens. As a proof of concept, we applied RAPPL-Seq to primary mouse lung and brain tissues (∼1 mg), human lung biopsies cryopreserved for more than 10 years, and surgically resected glioblastoma (GBM) and adjacent normal brain tissues from a patient. Notably, all human tissue samples weighed <1 mg, below the detection limit of the analytical balance **(Figure 4 and Supplemental Figure 4)**.

**Figure 4.**
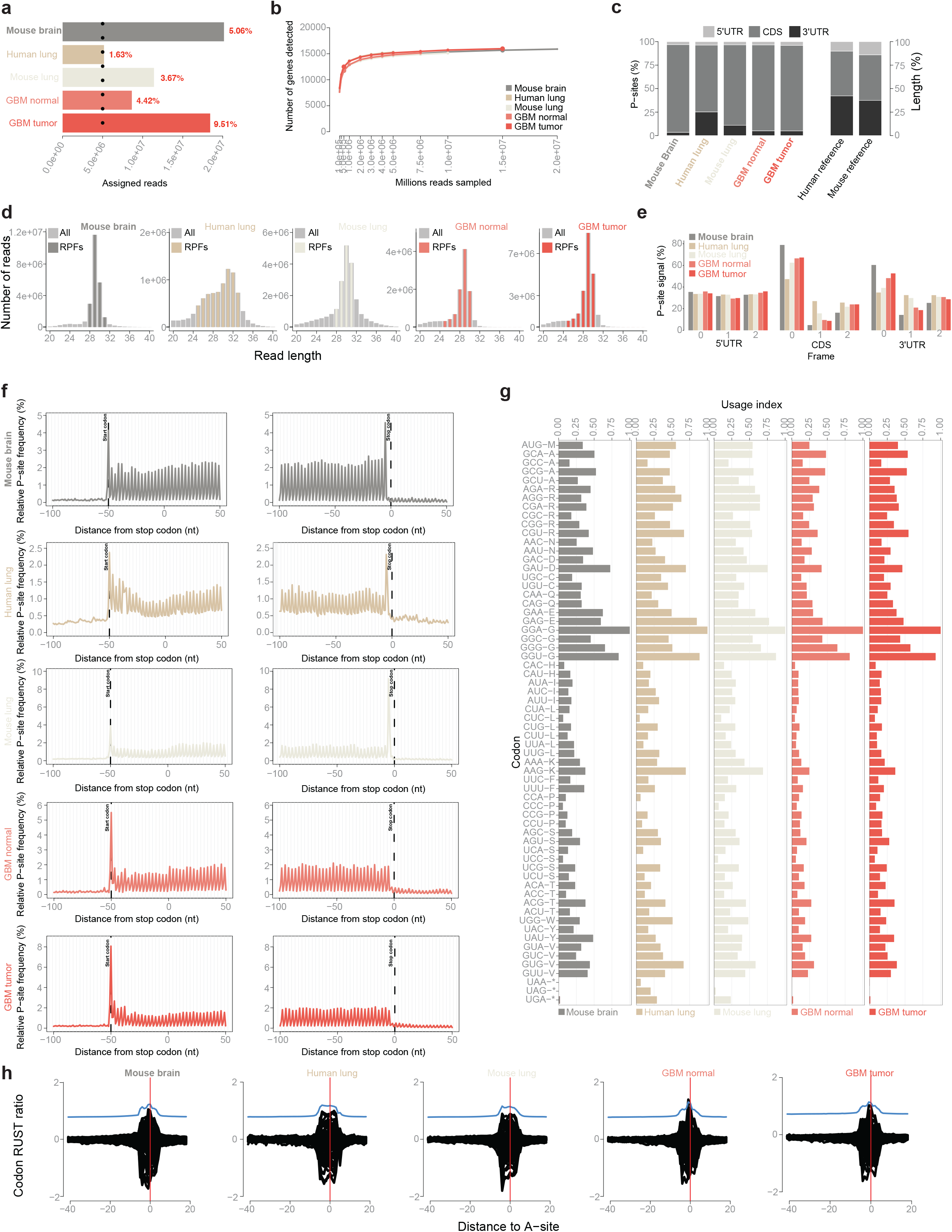
RAPPL-Seq is Broadly Applicable to Primary Tissues and Clinical Biospecimens. Tissue samples are colour-coded throughout the figure as follows: mouse brain (grey), human lung (tan), mouse lung (light beige), adjacent non-tumor brain tissue (“GBM normal”, salmon), and GBM tissue (“GBM tumor”, red). **(a)** Assigned RPF read counts and percentages out of total number of reads and **(b)** library complexity curves showing the number of genes detected across sampled sequencing depths for each sample. The dotted line in panel (a) indicates an approximate benchmark for differential translation analysis^29^. **(c)** Percentage of P-sites assigned to the 5’ UTR, CDS and 3’ UTR for each sample (left), compared to the expected distribution under random RNA fragmentation for mouse and human (right). **(d)** Read length distributions for all reads (grey) and RPFs (corresponding sample colours). **(e)** Distribution of P-sites among reading frames 0,1, and 2 within the 5’ UTR, CDS and 3’ UTR. **(f)** Metagene profiles showing relative P-site frequencies around annotated start codons (left) and stop codons (right), illustrating triplet periodicity within the CDS. **(g)** Codon usage profiles for each tissue sample. **(h)** Codon RUST ratios (log_2_) plotted relative to the A-site, indicated by the red vertical line. The corresponding Kullback–Leibler divergence is shown in blue.

Using RAPPL we achieved comparable RPF recovery from all tissue samples with exception of human lung biopsy **(Supplemental Figure 4a)**. The results of RAPPL-Seq on mouse and human tissues samples demonstrated distribution of total number and percentage of assigned RPF reads **(Figure 4a)** ranging from 5×10^6^ read counts and 1.63% of total reads in human lung biopsy, to 2×10^7^ and 5.06% of total reads for mouse brain sample. Mouse lung (1.1×10^7^ read counts and 3.67% of total reads), GBM patient resected normal (8.5×10^6^ and 4.42%) and tumor tissue (1.8×10^7^ and 9.51%) had similar proportion of assigned reads **(Figure 4a)** as compared to HEK293T cells **(Figure 1c)**. Moreover, the number of assigned RPF reads for all tissue samples reached the approximate read-count benchmark indicated for differential translation analysis (**Figure 4a; dotted line**). The overall number of detected genes was broadly comparable to that obtained with previously published methods and in HEK293T cells data **(Figure 1d and Figure 4b)**. Over 90% of assigned reads localized specifically to annotated protein-coding sequences (CDSs) with the largest proportion of the remaining assigned reads mapping to a subset of long non-coding RNAs (lncRNAs) **(Supplementary Figure 4b)**.

The high ribosome profiling quality was consistently observed across all tissue types examined. As expected, most P-site aligned RAPPL-Seq reads mapped predominantly to the CDS **(Figure 4c)**. RPF length distributions showed prominent peaks between 28 and 32 nt **(Figure 4d)**. A consistent preference for in-frame RPFs (frame 0) within the CDS **(Figure 4e)** and clear triplet periodicity in the metagene profiles **(Figure 4f)** further supported the quality of the captured footprints. Codon usage profiles were broadly similar, including between the paired GBM and non-tumor brain samples **(Figure 4g)**. Finally, we obtained similar codon RUST ratios for all tissue samples **(Figure 4h)**, which matched HEK293T samples **(Figure 1k)** and previous studies, thereby indicating absence of potential biases related to other sequence determinants. At the individual transcript level, RPF profiles showed high coverage of the housekeeping genes EEF2 and RPL8 **(Supplemental Figure 4c)**. Together, these results demonstrate that RAPPL-Seq is applicable to clinically relevant primary and biopsy samples, enabling translatome profiling throughout disease progression.

### RAPPL-Seq Efficiently Captures Translatomes from Unaffected Brain Tissue and GBM Tumor Biopsies

Differences in mRNA translation between tissues, as well as disease associated translational reprogramming, cannot be directly quantified by standard mRNA-seq^15–18^. Characterizing these changes in clinical samples has been hindered by the scarcity of biopsy material. Building on the performance of RAPPL-Seq with low input tissue samples, we next examined paired RNA-seq and RAPPL-Seq from mouse and human tissues (i.e. same as **Figure 4** and **Supplementary Figure 4)**.

To assess whether RAPPL-Seq captures tissue specific translation markers, we first examined RPFs reads distributions for the brain specific Synaptosomal-Associated Protein 25 (SNAP25)^53^ and myelin-associated glycoprotein (MAG)^54^, as well as pulmonary specific gene markers Surfactant Protein D gene (SFTPD)^55^, Plasmalemmal vesicle-associated protein (PLVAP)^56^, Napsin A (NAPSA)^57^ and Claudin-18.1 (CLDN18)^58^ **(Figure 5a-d)**. Notably, RAPPL-Seq accurately captured the translation of selective brain markers SNAP25 and MAG in the mouse and GBM patient brain samples, with minimal signal in mouse and human lung samples **(Figure 5a and 5c)**, confirming the preservation of brain tissue-type signatures. In parallel to brain specific gene RPF distributions, pulmonary gene markers SFTPD, PLVAP, NAPSA and CLDN18 showed robust translation and RPFs distribution in datasets obtained from mouse lung and human lung biopsy samples with notable absence of RPFs in the mouse or human brain tissue samples **(Figure 5b and 5d)**. These tissue specific gene RPFs distributions are in sharp contrast to previously shown uniform RPF read distribution in common and housekeeping EEF2 and RPL8 genes present in all tissue samples **(Supplemental Figure 4c)**, further demonstrating that RAPPL-Seq efficiently captures tissue-specific translatomes.

**Figure 5.**
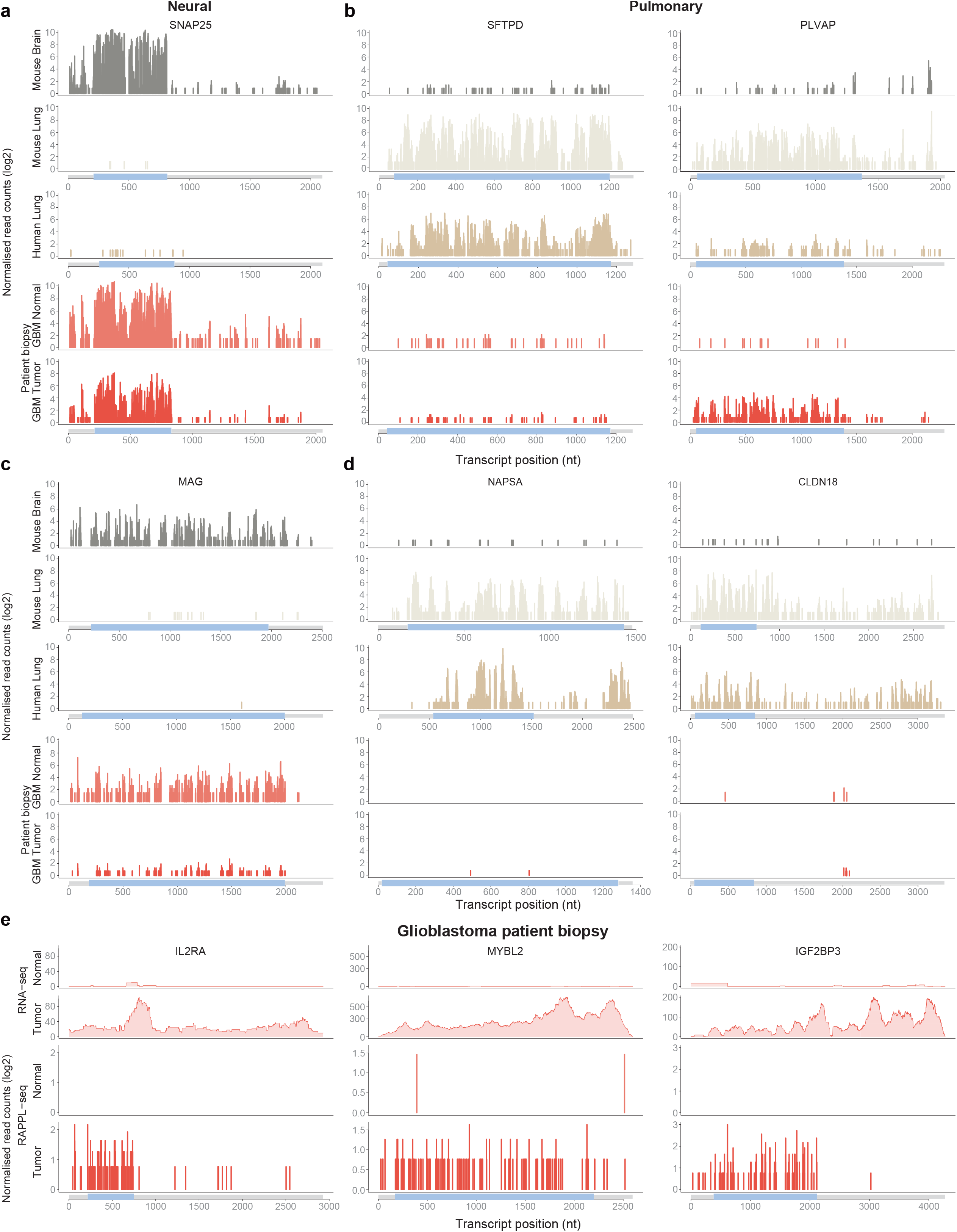
RAPPL-Seq Reveals Tissue- and Tumor-Specific mRNA Translation Profiles. Tissue samples are colour-coded throughout the figure as follows: mouse brain (grey), human lung (tan), mouse lung (light beige), adjacent non-tumor brain tissue (“GBM normal”; salmon), and GBM tissue (“GBM tumor”; red). **(a-d)** Representative RAPPL-Seq coverage profiles shown as log_2_ TMM-normalised P-site counts along transcript position. **(a)** SNAP25, a neuronal marker gene. **(b)** SFTPD and PLVAP, lung-associated marker genes. **(c)** MAG, a neural-associated transcript. **(d)** NAPSA and CLDN18, lung specific markers. Transcript models are shown below each set of profiles. **(e)** Representative RNA-seq (raw counts, top) and RAPPL-Seq profiles (log_2_ TMM-normalized) of IL2RA, MYBL2, and IGF2BP3 in the paired GBM tumor and adjacent non-tumor brain samples, illustrating tumor-associated differences in mRNA translation.

Interestingly, analysis of patient GBM tumor sample, and not adjacent normal brain tissue, indicated reduced distribution and less RPFs in the brain specific MAG transcript **(Figure 5c)**. In parallel, the resected patient GBM tumor sample had high number and CDS specific distribution of RPFs in lung-specific PLVAP transcript, which was absent in adjacent normal brain tissue **(Figure 5b).** This data argues for potential loss of tissue-specific translational control in tumor cells. To get further insight into possible protein synthesis differences between tumor and adjacent normal tissues from the GBM patient sample we focused on RAPPL-Seq data from the alpha subunit of the high-affinity Interleukin-2 receptor (IL2RA), MYB proto-oncogene like 2 (MYBL2) and Insulin-like growth factor 2 mRNA-binding protein 3 (IGF2BP3) **(Figure 5e).** All three transcripts had considerable and CDS-specific distribution RPF reads in the tumor sample but with minimal signal in adjacent normal tissue **(Figure 5e).** IL2RA is associated with T-cells infiltrating GBM tumors^59^, while MYBL2 and IGF2BP3 are well described glioblastoma-specific and oncogenic drivers^60–62^. Finally, we found multiple other transcripts with a significant amount of RPFs in CDSs that were present mostly in tumor and not adjacent normal brain tissue from GBM patient **(Supplemental Figure 5a)**. These included GBM-cell associated and known tumor cell survival and progression genes like Protein Kinase-Membrane Associated Tyrosine/Threonine 1 (PKMYT1)^63,64^, Survivin (BIRC5)^65,66^ and NADPH oxidase 4 (NOX4)^67^, as well as a known biomarker for GBM and gliomas - AKNA Domain Containing 1 (AKNAD1) transcript^68^. We also found an increase in RPFs in CDSs of multiple HOX transcription factors (HOXA2, HOXA4, HOXA7, HOXB4, HOXB7, HOXC4, HOXD8), suggesting known activation and translation of these genes in GBMs^69–71^. Together, our findings demonstrate that RAPPL-Seq enables the robust and systematic profiling of translation in different tissues and distinct tissue states (i.e., healthy vs. diseased) from scarce human biopsy samples.

## DISCUSSION

Ribosome profiling enables transcriptome-wide monitoring of translation *in vivo* under normal, stress, and disease conditions through high-throughput sequencing, offering critical insights into the molecular underpinnings of translational regulation. A major limitation of traditional ribosome profiling lies in the lengthy protocols and non-efficient isolation of RPFs. These procedures typically require expensive equipment, such as ultracentrifuges, where ribosomes and ribosome-associated RPFs are separated by sucrose cushions or gradients followed by RNA precipitation. The sucrose cushions or gradients impact the yield and recovery of ribosomes and RPFs as they are inherently inefficient, often resulting in substantial material loss (∼80-95%)^24^. These approaches thereby restrict the utility of ribosome profiling in settings with limited input material, such as clinical biopsies or primary tissues, and reduce sensitivity of these approaches for lowly expressed and translated RNAs.

To overcome these limitations, we developed RAPPL-Seq, an adaptation of RAPPL^24^, which achieves significantly higher recovery efficiency (>80%) within a short processing time (<3 h) and eliminates the need for specialized equipment such as ultracentrifuges and gradient fractionators. This method enables faster ribosome profiling from as few as 200,000 cells and tissue samples, substantially broadening its applicability to low-input and clinically relevant samples. Notably, while a recent study introduced a cell-sorting-based single-cell ribosome profiling technique^72^, that approach is incompatible with cryopreserved clinical specimens due to poor cell sorting recovery after thawing the samples. Furthermore, it remains less suited for low-input samples due to RPF loss during cell sorting and high rRNA contamination resulting from the lack of an adaptable rRNA depletion workflow^72^. In contrast, RAPPL-Seq is a fast and resilient method that efficiently, and with high-reproducibility, captures RPFs from different cell numbers (100,000 to 1,000,000). However, at lower cell threshold, standard RNase I digestion conditions may become overly stringent, leading to RPF over-digestion - a phenomenon documented in other low-input applications^73^. For samples containing 100,000 cells or fewer, optimization of digestion conditions will be necessary, including adjusting RNase I concentration and incubation time or employing alternative nucleases with lower degradative activity, such as P1 Nuclease^74^ and RNase-T1^75^.

RAPPL-Seq demonstrates adaptability to multiple translation inhibitors, making it well suited for comparative analyses of compounds that arrest translation at distinct stages^1,4,39,40,42^. By comparing ribosome profiles from liquid nitrogen snap-frozen samples with those obtained following treatment with specific inhibitors, this approach enables characterization of the precise inhibitory mechanisms of different inhibitors in the same fashion as other ribosome profiling methods but with better gene coverage and sequencing depth. Furthermore, RAPPL-Seq is broadly applicable to clinically relevant samples, such as primary tissues, biopsies and surgical tumor resections, which are often limited by low cell numbers and remain challenging for traditional ribosome profiling. This capability allows for the monitoring of translatome profiles directly in patient tissue biospecimens, providing critical insights into tissue specific translational regulation or protein synthesis during disease progression.

Using RAPPL-Seq we were able to capture tissue specific translation profiles from mouse brain and lungs, as well as from human lung biopsies that have been frozen for ∼10 years. These samples also helped indicate differences in the expression of tissue specific genes in the tumor and adjacent normal tissue surgically removed from a GBM patient, mainly indicating increase in brain non-specific transcript translation in tumor samples. Ultimately, by capturing the active translatome from GBM patient tumor resection, we were able to investigate tumor-specific protein synthesis that revealed activation and robust translation of multiple GBM-and T-cell infiltrating transcripts in tumor but not normal tissue. These analyses remain largely undetectable by standard transcriptomic or classical ribosome profiling analyses^16,76–79^.

In summary, RAPPL-Seq provides a fast, robust, and accessible platform for ribosome profiling from limited input, delivering higher sensitivity and a shorter turnaround time than existing methods. Its ability to uncover translational regulation in clinically relevant samples, such as primary tissues and biopsy specimens, positions RAPPL-Seq as a powerful tool for mapping translatome dynamics during disease progression. By bridging the gap between current transcriptomic and proteomic profiling, this approach reveals a new and crucial layer of the regulatory insight of disease pathogenesis. Ultimately, RAPPL-Seq has the potential to transform disease diagnosis and accelerate the discovery of novel drug targets, benefiting both basic research and translational medicine.

## METHODS

### Cell Culture and Harvest

#### (i) HEK293T

HEK293T cells were cultured in StableCell DMEM-high glucose (Sigma-Aldrich) supplemented with 10% fetal bovine serum (Gibco) under 5% CO₂ until ∼90% confluency. Two 10-cm plates were prepared: one plate was harvested and counted using an automated cell counter (Countess 3, Thermo Fisher Scientific), while the other was washed twice with cold PBS and snap-frozen by direct addition of LN_2_. For translation inhibitor treatments, cells were incubated in media containing either 100 µg/mL cycloheximide (CHX) or 2 µg/mL harringtonine (Harr) at 37°C for 10 min, followed by two cold PBS washes. Cells were then lysed in 1 mL lysis buffer (10 mM Tris-HCl, pH 7.5; 100 mM NaCl; 5 mM MgCl_2_; 1% Triton X-100; 0.5% sodium deoxycholate (NaDoc); and 0.5 mM dithiothreitol (DTT)) for 10 min on ice, scraped, and centrifuged at 14,000 rpm for 10 min at 4 °C. Supernatants were collected and aliquoted according to the desired cell numbers determined from the counted plate, and volumes were adjusted to 400 µL with lysis buffer for either sucrose gradient fractionation or RAPPL-Seq. For CHX- and Harr-treated samples, the respective inhibitor (100 µg/mL CHX or 2 µg/mL Harr) was also included in the lysis buffer.

#### (ii) Mouse primary tissues and human tissue biopsies (low-input material)

The mouse lung and brain tissues were harvested from adult C57BL/6 mice (The Jackson Laboratory: #000664). Mice were euthanized by CO_2_ asphyxiation for lung and brain collection, followed by immediate dissection. Freshly isolated tissues were snap-frozen in LN_2_ and stored at −80°C. The archived snap-frozen human lung biopsies from healthy donors were obtained from Dr. Zhiyu Dai from the Pulmonary Hypertension Breakthrough Initiative (PHBI)^80^ \. The archived snap-frozen human brain tumor (contrast-enhancing portion of the tumor) and matched adjacent normal brain (not involved brain by imaging) from a patient with histologically confirmed glioblastoma were obtained from Dr. Dimitrios Mathios through the CNS Tumor Bank at the Brain Tumor Center at Washington University in St. Louis. For tissue homogenization, frozen mouse tissue (1-2 mg) or human tissue (all the tissue biopsies; <1 mg) was minced with scissors and homogenized in 600 µL of CHX-containing cell lysis buffer using 1.5 mm zirconium beads (Zymo Research). Homogenization was performed using a BeadBug 6 homogenizer (Benchmark Scientific) at 4,000 rpm for 2-3 cycles (on/off: 30s/30s) at 4°C. Lysates were centrifuged at maximum speed for 5 min at 4°C, and 400 µL of the supernatant was transferred to new 1.5 mL microcentrifuge tubes as cell lysates.

### RPF Generation

RPF generation from the cell lysate was adapted from Sharma et al^35^. Specifically, cell lysates were digested with 900 U RNase I (Ambion) per 10 A_260_ units of RNA, after subtracting background A_260_ from the lysis buffer (900 × (A_260_lysate - A_260_buffer) / 10), at 22 °C for 1 h on a Thermomixer (Eppendorf) with shaking at 800 rpm. RNase I activity was subsequently quenched by adding 150 U SUPERase·In™ RNase Inhibitor (Invitrogen) per 10 A_260_ units of RNA.

### RAPPL-Seq

#### (i) RPF isolation

30 µL of poly-lysine (polyK) magnetic beads (MacLab) were washed twice with lysis buffer (supplemented with CHX or Harr for inhibitor-treated samples) and incubated with RPF samples at 4 °C for 30 min on a rotator. Beads were then washed three times with 500 µL washing buffer (20 mM Tris-HCl, pH 7.5; 5 mM MgCl_2_; 100 mM NH_4_Cl; and 1 mM DTT; 150 mM NaCl) for 1 min each time. RPFs were eluted twice from beads, each time with 14 µL elution buffer (20 mM Tris-HCl, pH 7.5; 5 mM MgCl₂; 100 mM NH₄Cl; 1 mM DTT; and 2 mg/mL poly-D-glutamic acid (polyE; Sigma #4033)) supplemented with 0.5 µL SUPERase·In™ RNase Inhibitor (Invitrogen), at 37 °C for 10 min on a Thermomixer (Eppendorf) with shaking at 1300 rpm. Eluted RPFs were pooled, mixed with 28 µL RNA Gel Loading Dye (2X) (Thermo Fisher Scientific), heated at 70 °C for 5 min, and loaded into two lanes of a 15% TBE-Urea gel (Mini-PROTEAN®, 10-well, 30 µL; Bio-Rad) for electrophoresis at 180V for 40 minutes. After staining the gel with SYBR™ Gold (ThermoFisher Scientific) at room temperature for 5 min, RPFs of 25-35 nt were excised using RNA size markers of 25 nt (5’-GACUCUUAGCGGUGGAUCACUCGGC-3’) and 35 nt (5’-GACUCUUAGCGGUGGAUCACUCGGCUCGUGCGUCG-3’). Two gel slices containing the RPFs from the same sample were pooled and purified using the ZR Small-RNA PAGE Recovery Kit (Zymo Research) according to the manufacturer’s protocol.

#### (ii) Library generation and sequencing

Human or mouse rRNAs and tRNAs were depleted from RPFs using the RiboMinus™ Eukaryote Kit v2 (Thermo Fisher Scientific) with an additional spike-in of 1 µL Human or Mouse tRNA (Ribo-Seq) riboPOOL Probes (Galen Molecular), respectively. The rRNA/tRNA-depleted RPF libraries were prepared using the LACEseq Library Prep Kit (Immagina) and Illumina iUDI index primers (Immagina) according to the manufacturer’s protocol. For low-input samples, including the 200,000-cell HEK293T aliquots, mouse primary tissues, and human clinical biopsies, an additional size-selection step was integrated. Following the standard LACEseq preparation, libraries were resolved on a 4% E-Gel EX agarose gel (Invitrogen: G401001) to specifically isolate and enrich for the ∼200 bp target RPF library fragments. The final prepared libraries were sequenced on an Illumina NovaSeq X Plus platform utilizing a single-end 150-cycle (SE150) run configuration, targeting a sequencing depth of approximately 200 million reads per sample.

### mRNA sequencing (mRNA-seq)

A fraction (1-5%) of the cell lysate collected for RAPPL-Seq was reserved for mRNA-seq analysis. To reduce detergent concentration, the lysate was diluted 10-fold in water, followed by RNA purification using the RNA Clean & Concentrator Kit with on-column DNase I digestion (Zymo Research). mRNA libraries were prepared using the NEBNext® Ultra™ II RNA Library Prep Kit (NEB #E7775) coupled with the NEBNext Poly(A) mRNA Magnetic Isolation Module (NEB #E4790), following the manufacturer’s protocol. For the final PCR amplification, commercial primers were replaced with custom adapters (i5 primer: AATGATACGGCGACCACCGAGATCTACAC-N10(index)-ACACTCTTTCCCTACACGACGCTC; i7 primer: CAAGCAGAAGACGGCATACGAGAT-N10(index)-GTGACTGGAGTTCAGACGTGT) to enable unique dual indexing and minimize index hopping. Libraries were sequenced on an Illumina NovaSeq X Plus platform using a single-end 150-cycle (SE150) run configuration, targeting 50 million reads per sample.

### RAPPL-seq and RNA-seq analysis

Raw RAPPL-seq reads were processed according to the LACEseq BioIT guidelines (v3; Immagina). The 3′ adapter (TCTCCTTGCATAATCACCAACC) was removed using Cutadapt^81^ (v1.18), with untrimmed reads and reads shorter than 20 nt were discarded (--discard-untrimmed -m 20 --nextseq-trim=20). UMIs were extracted using UMI-tools^82^ (--bc-pattern=’^(?P<umi_1>.{4}).+(?P<umi_2>.{4})$’ --extract-method=regex). The thymine preceding the ribosome-protected fragment was subsequently removed using Cutadapt (-g ^T --discard-untrimmed). Reads matching rRNA sequences from the SILVA database^83^ were removed using BBMap^84^. The remaining reads were aligned to the hg38 or mm39 genome using Bowtie^85^ (v1.2.2) (-a -m 1 --best --strata -n 2 -l 20). Reads that aligned uniquely or remained unaligned were subsequently mapped to the corresponding RefSeq^86^ transcript sequences (human: GCF_000001405.40; mouse: GCF_000001635.27) using Bowtie (-a --best --strata -n 2 -l 20). PCR duplicates were removed using UMI-tools based on alignment coordinates and UMI sequence. RPF length ranges indicated in the corresponding figures were retained. P-site assignment, metagene profiling and quality-control analyses were performed using riboWaltz^87^ with the same RefSeq annotations. Ribo-seq Unit Step Transformation (RUST)^38^ analysis was used to identify sequence-specific biases at ribosome-footprint boundaries. For downstream gene-level analyses, one transcript was randomly selected as the representative of each gene within each dataset to avoid isoform redundancy. Read counts were normalised between samples using the trimmed mean of M-values (TMM)^88^ method implemented in edgeR^89^.

For gene-biotype analysis, reads were additionally aligned to the genome using HISAT2^90^ (v2.0.4). Primary alignments with NH:i:1 were considered uniquely mapped and used to filter the transcriptome alignments. Gene biotypes were assigned using GENCODE^91^ (human: v50, mouse: vM39). Reads mapping to multiple non-coding biotypes were classified as ambiguous_non_protein, whereas any read mapping to a protein-coding gene was classified as protein_coding.

RNA-seq reads were preprocessed using BBMap (v36.59) (k=13 ktrim=n useshortkmers=t mink=5 qtrim=t trimq=10 minlength=25). Processed reads were aligned to the hg38 reference genome using HISAT2 (v2.0.4) with default settings.

### RPF isolation with sucrose gradient

Sucrose gradient-based RPF isolation was performed as described in Erath et al.^24^, with modifications to buffer compositions and sample processing steps as detailed in this study. Equal numbers of HEK293T cells were plated on 10-cm dishes and, 24 h later, treated with 100 μg/mL CHX for 15 min prior to harvesting. Cell lysates were prepared as for RAPPL-Seq with RNase I treatment. One lysate was processed using the RAPPL-Seq procedure, while the other was subjected to sucrose gradient isolation. Specifically, lysates were layered over a 5-50% sucrose gradient (20 mM HEPES, 200 mM KCl, 10 mM MgCl₂, 1 mM DTT, 100 μg/mL CHX) prepared with a Gradient Master 108 (BioComp) and centrifuged at 209,490 × g (35,000 rpm) for 2.5 h at 4 °C in an SW41 Ti swinging-bucket rotor (Beckman Coulter). Ribosome profiles were resolved using a Brandel gradient fractionator with continuous absorbance monitoring at 254 nm (Brandel UA-6). Fractions were pooled and divided into two portions: one processed with RAPPL and the other with standard RNA precipitation^21^. For RAPPL samples, RPFs were eluted with polyE elution buffer and purified using the RNA Clean & Concentrator Kit (Zymo Research) according to the manufacturer’s protocol. RNA recovery efficiency was calculated by back-tracking sample splitting at each step.

### Western Blotting

500,000 HEK293T cells were cultured, harvested, and processed as described in the RAPPL-Seq workflow with or without CHX treatment and with or without RNase I digestion prior to RPF purification. After elution from poly-lysine magnetic beads, samples were supplemented with XT loading buffer, boiled for 5 mins at 95°C, and cooled on ice. Samples were loaded onto pre-cast gels (4–12% Criterion™ XT Bis-Tris Protein Gels, Bio-Rad Laboratories, #3450124 and #4561085) with protein markers (Bio-Rad Laboratories #1610374 and #1610373 and ELISHA’s markers). Samples were run for one hour at 180V. Gels were transferred to PVDF membrane using a semi-dry transfer systems (Bio-Rad Laboratories #1703940 and #1704150) for 17-30 mins at 25V. Membranes were blocked in 1X PBS + 0.1% Tween 20 + 5% dry milk for 1 hour at RT. Membranes were blotted in blocking reagent (above) 4℃ overnight. Primary antibodies (1:1000) that were not HRP-conjugated were blotted as above with secondary antibodies (1:5000) that were HRP conjugated. Blots were visualized using SuperSignal™ West Pico PLUS Chemiluminescent Substrate (Thermo Fisher Scientific #34580) and SuperSignal™ West Femto Maximum Sensitivity Substrate (Thermo Fisher Scientific # 34096) with subsequent imaging using a Bio-Rad Laboratories imaging system.

## ACKNOWLEDGEMENTS

Next-generation sequencing was done at the Genome Technology Access Center (GTAC) at Washington University in St. Louis. This work was funded by NIH R00HG011475 and NIH R35GM165632 to C.-K.C., NIH R35GM167503, NCI U01CA305187, Chen-Zuckerberg Neurodegeneration Initiative, and the Siteman Cancer Center Investment Award to S.D. We would like to acknowledge the CNS Tumor Bank of the Brain Tumor Center at Washington University in St Louis for providing high-quality human specimens for this work. IT is supported by the Canadian Institutes of Health Research (CIHR) (PJT-581421), DNA to RNA (D2R) Initiative at McGill University, funded by the Canada First Research Excellence Fund (D2R-FDN2-A2-95), and the Canada Research Chair in Regulation of mRNA Translation and Metabolism. O.L. is supported by grants from Swedish Research Council (2025-03216), Swedish Cancer Society (25 4545 Pj), the cancer research funds of Radiumhemmet (231263).

## AUTHOR CONTRIBUTIONS

A.P., K.J.S, C.-K.C, and S.D. led the project. A.P. and C.L. conducted experiments. K.J.S, S.D., and C.-K.C analyzed the data. J.E. and S.P.D. carried out Western blotting and sucrose gradient experiments. I.T., K.J.S, and O.L. assisted the computational analysis. Z.D. collected and processed the human lung biopsies. X.H. harvested and processed mouse primary tissues. J.H. and D.M. collected and processed the human GBM and matched adjacent normal brain biopsies. C.-K.C. and S.D. conceived and oversaw the overall project. A.P., K.J.S, I.T., S.D., and C.-K.C. wrote the manuscript with input and critical revisions from all authors.

## DECLARATION OF INTERESTS

Washington University in St. Louis has filed International Patent Application No.: PCT/US2025/046232 “Rapid Isolation of Ribosomes and Associated Molecules” based on this work, and J.E., S.P.D., C.-K.C., and S.D. are named as co-inventors.

## SUPPLEMENTAL FIGURE LEGENDS

**Supplemental Figure 1. RAPPL-Seq enables Rapid, Efficient, and Consistent Capture of RPFs. (a)** Western blotting shows efficient ribosome recovery with RAPPL. uS9: small ribosomal subunit protein; Inp: Input; FT: flow through after RAPPL binding; W1: flow through in the first wash solution; RAPPL: elution from RAPPL. CHX: cycloheximide treatment. **(b)** Schematic of the isolation of ribosomes and generation of RPFs using sucrose gradient (SG) and RAPPL. Data are shown as mean ± SEM (% of starting material or % of SG denotes) of 3 independent replicates. **(c)** Read distribution across preprocessing categories, showing the fractions removed during adaptor filtering, rRNA depletion, and mapping, and the final proportion retained as assigned RPF reads.

**Supplemental Figure 2. RAPPL-Seq Enables Analysis of Translation Inhibitors Acting at Different Stages.** The same colour coding identifies the three conditions throughout the figure: harringtonine (Harr, brown) and cycloheximide (CHX, teal) pretreatments, and flash-freezing in liquid nitrogen (LN_2,_ blue). **(a)** Distribution of P-sites among reading frames 0, 1, and 2 within the 5′ UTR, CDS, and 3′ UTR. **(b)** Codon usage profiles for each condition. **(c-f)** Representative RNA-seq coverage profiles (raw counts, top) and RAPPL-Seq profiles (log_2_ TMM-normalised P-site counts) under the three conditions for RNF10 **(c)**, RAB6A **(d)**, CCDC124 **(e)**, and ACTB **(f)**.

**Supplemental Figure 3. RAPPL-Seq Enables Ribosome Profiling of Transcripts Across a Range of Expression Levels.** The same colour coding identifies the four input amounts throughout the figure: 1000K (dark blue), 500K (slate blue), 200K (blue grey), and 100K (light grey) HEK293T cells. **(a)** Distribution of transcript RPF abundance in the 1000K sample, with representative genes highlighted to indicate high-, medium-, and low-abundance groups. **(b-d)** Representative RNA-seq profiles (raw counts, top) and RAPPL-Seq profiles (log_2_ TMM-normalised P-site counts) of highly expressed transcripts DDX1, MCM6, and CCT3 **(b)**, moderately expressed transcripts DDX59, UTP4, and RBM14 **(c),** and lowly expressed transcripts PNISR, BRAF, and KMT2A **(d).** Transcript models are shown below each set of profiles.

**Supplemental Figure 4. RAPPL-Seq Profiling Across Primary Tissues and Clinical Biospecimens. (a)** Read distribution across preprocessing categories, showing the fractions removed during adaptor filtering, rRNA depletion, and mapping, and the final proportion retained as assigned RPF reads for mouse brain, human lung biopsy, mouse lung, adjacent non-tumor brain tissue (“GBM normal”), and GBM tissue (“GBM tumor”). **(b)** Distribution of RNA biotypes among RAPPL-Seq reads mapped to GENCODE (human: v50, mouse: vM39) transcripts. **(c)** Representative RAPPL-Seq coverage profiles (log₂ TMM-normalised P-site counts) for the housekeeping genes RPL8 and EEF2 across the five tissue samples. Transcript models are shown below each set of profiles.

**Supplemental Figure 5. RAPPL-Seq Reveals Tumor-Associated Translation Profiles in GBM**. Samples are shown as adjacent non-tumor brain tissue (“Normal”) and GBM tumor tissue (“Tumor”). **(a)** Representative RNA-seq profiles (raw counts, top) and RAPPL-Seq profiles (log₂ TMM-normalized P-site counts) for AKNAD1, PKMYT1, BIRC5, NOX4, OTX2, HOXC4, HOXA7, HOXB7, HOXD8, HOXA4, HOXB4, and HOXA2 in the paired GBM tumor and adjacent non-tumor brain samples. Transcript models are shown below each set of profiles.

