## Supplemental Figures for "RAPPL-Seq Enables Rapid, High-Sensitivity Ribosome Profiling of Scarce Samples"

Supplemental Figure 1

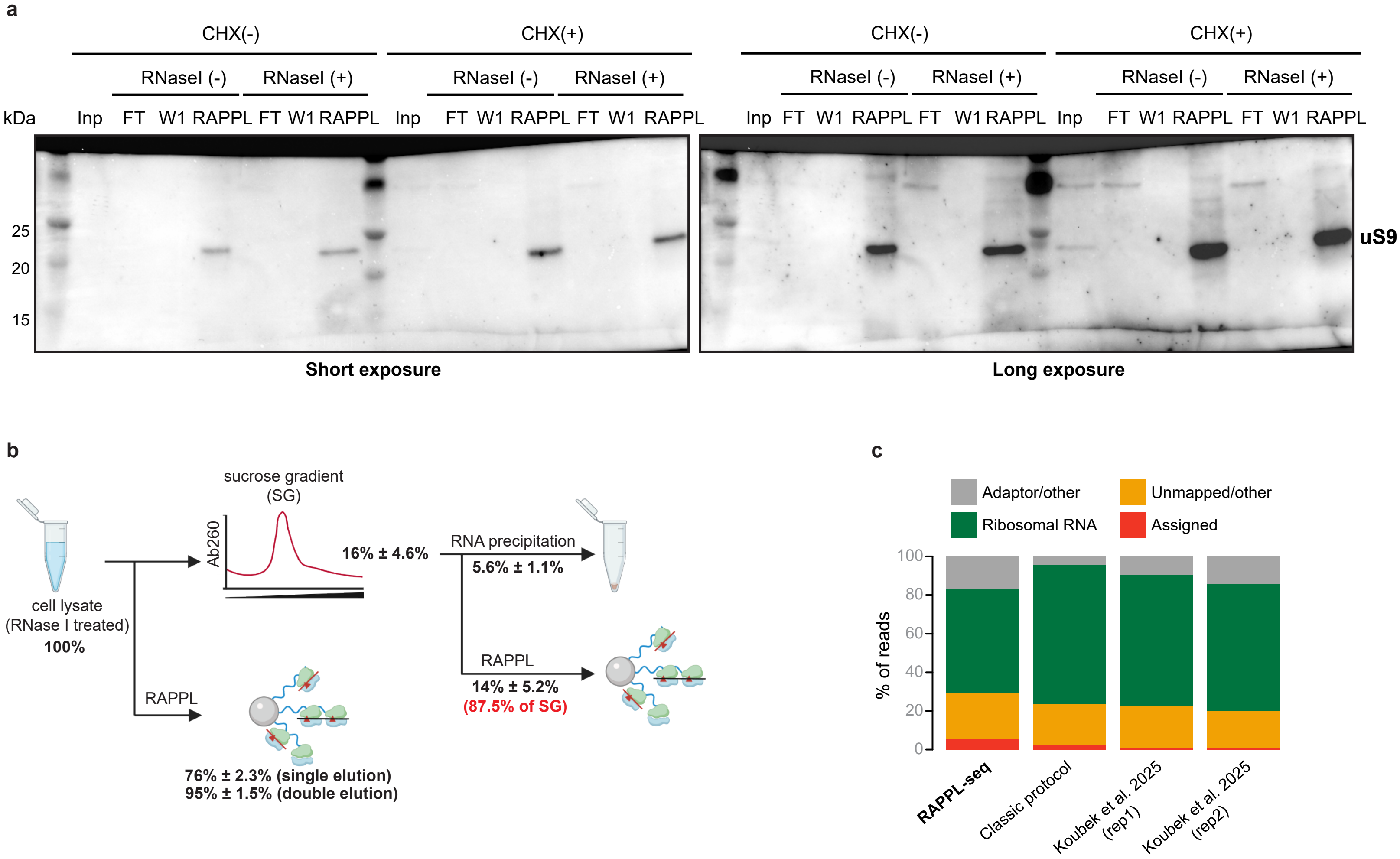

**Supplemental Figure 2**

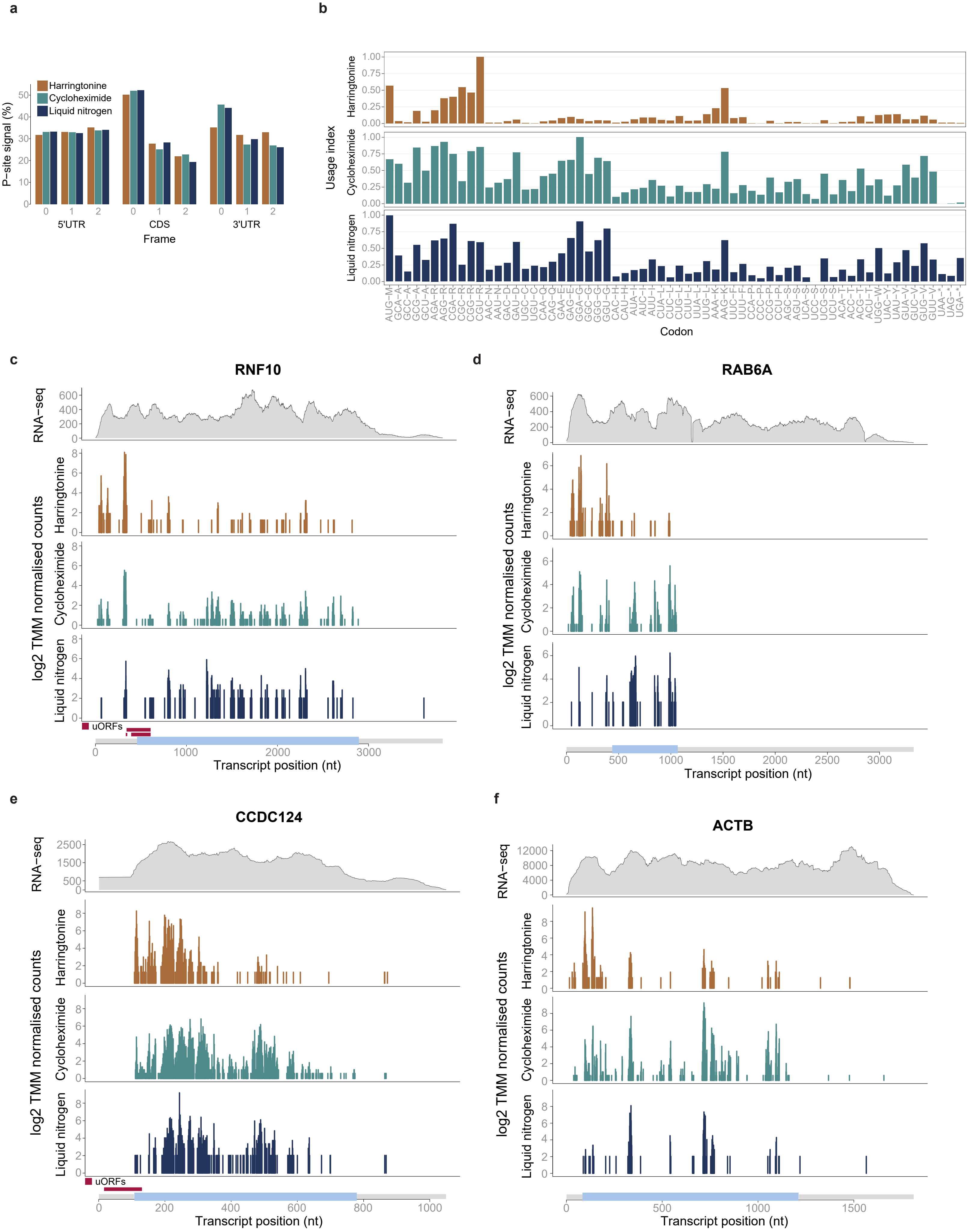

**Supplemental Figure 3**

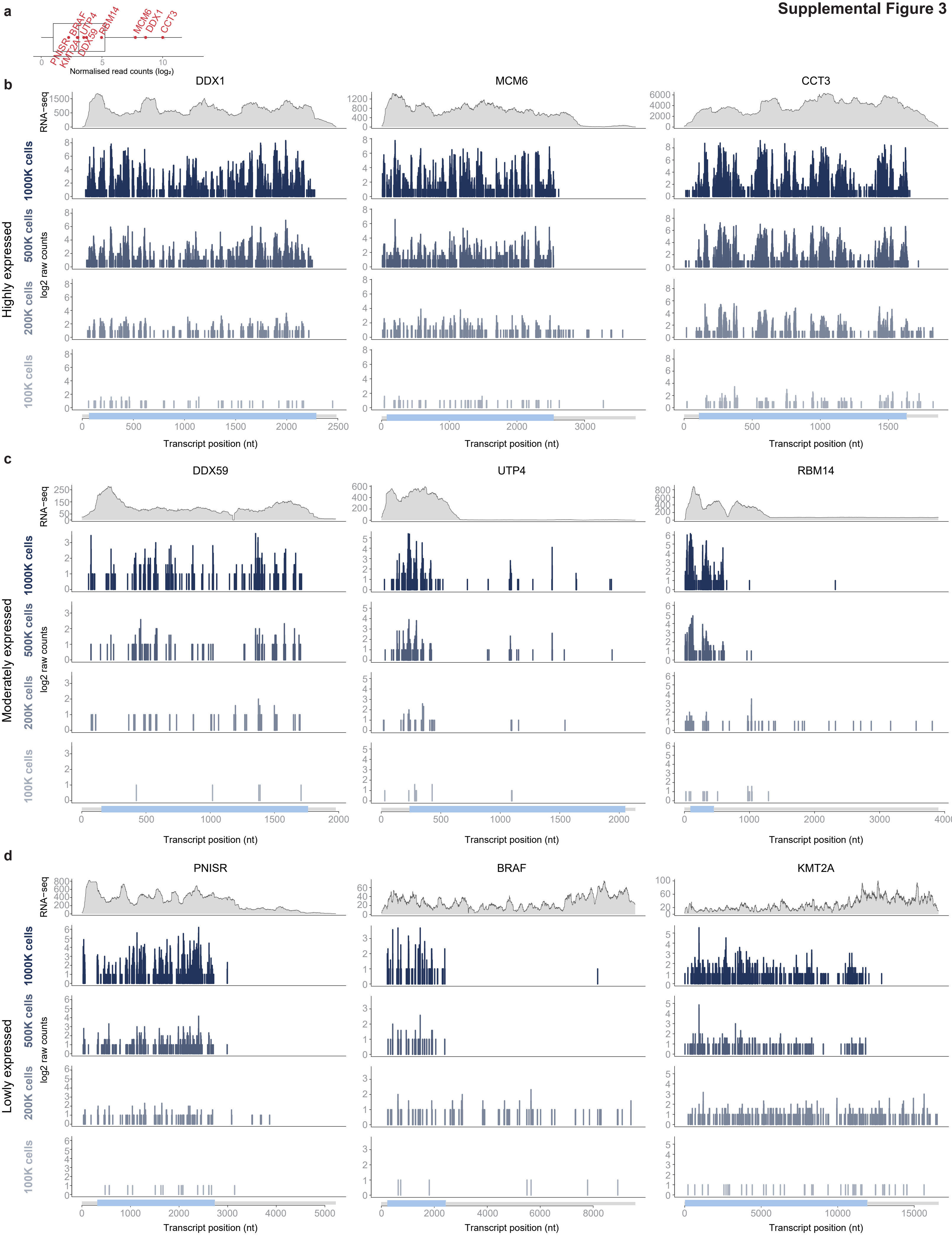

**Supplemental Figure 4**

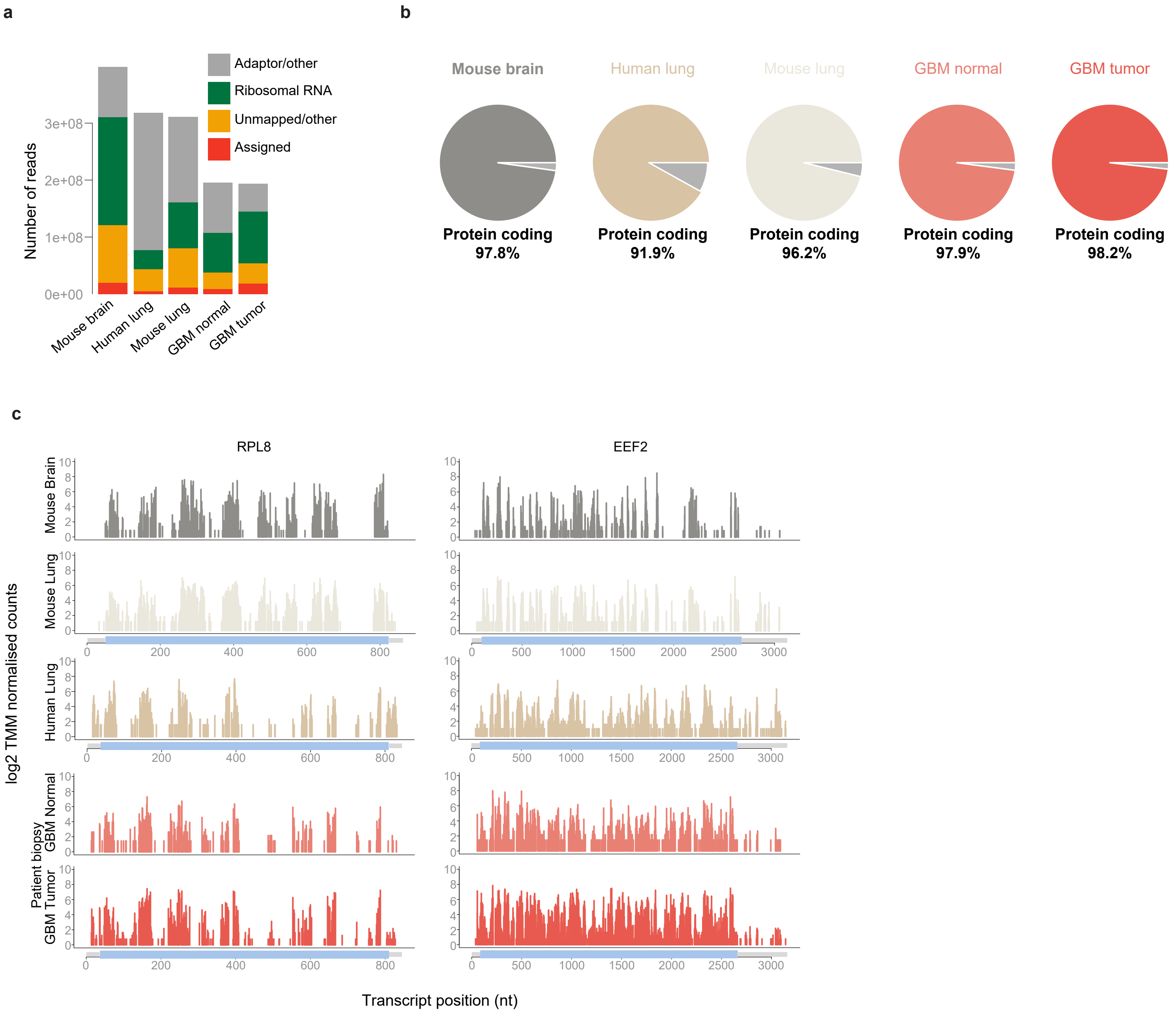

**Glioblastoma patient biopsy**

**Supplemental Figure 5**

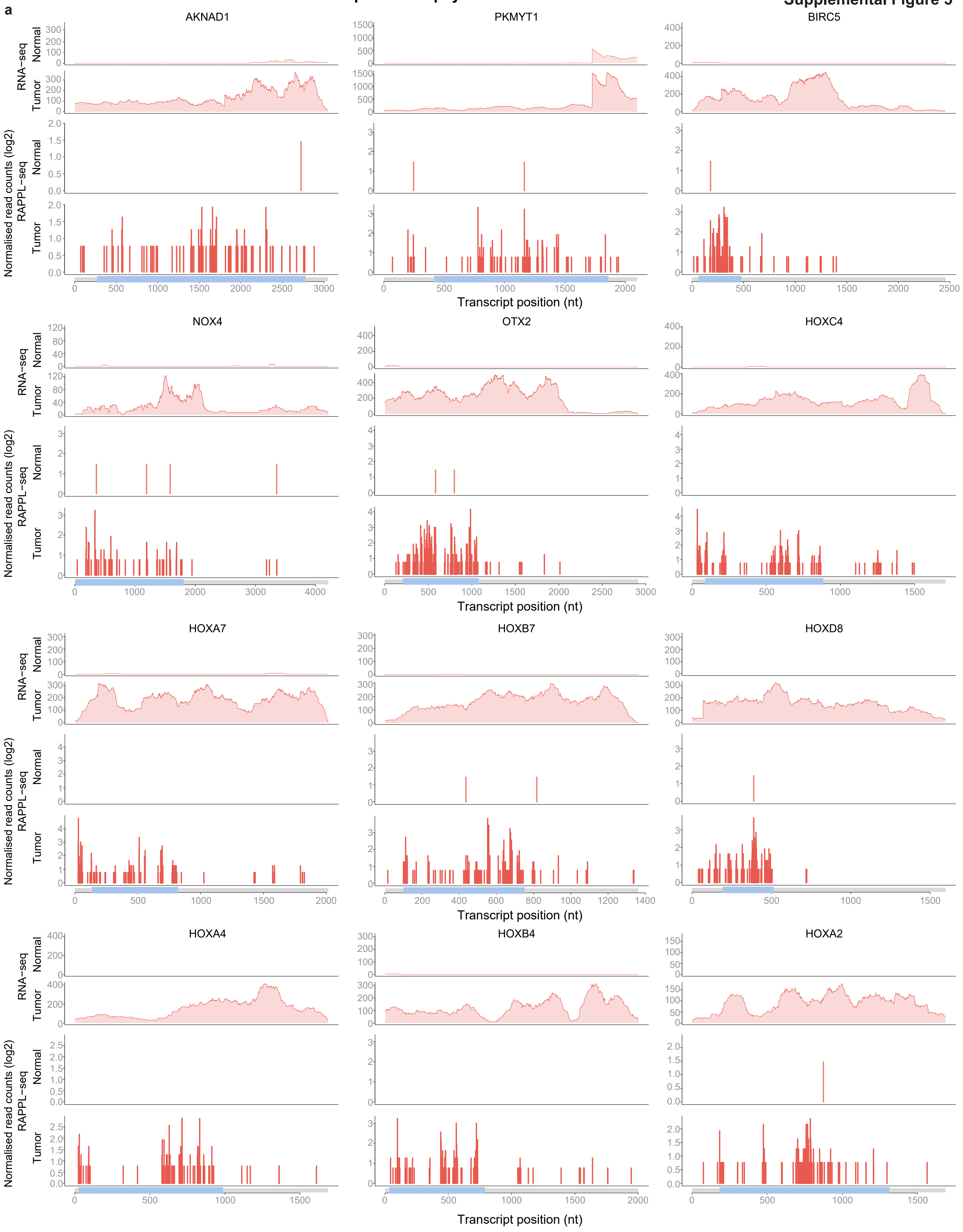
